# The Trk fused gene-product (Tfg) is part of a 600-700kDa CARMA1 complex

**DOI:** 10.1101/857342

**Authors:** Markus Grohmann, Tobit Steinmetz, Hans-Martin Jäck, Dirk Mielenz

## Abstract

B cell receptor (BCR) mediated activation of nuclear factor κB (NF-κB) is key to humoral immunity. CARMA1 (CARD11) is essential for BCR mediated NF-κB activation by interacting with Bcl10 and MALT1. Besides these two main players, few interaction partners of the CARMA1 complex are known. Here we identified new interaction partners of CARMA1. We generated two rabbit antisera against mouse CARMA1 to immunopurify endogenous CARMA1 from lysates of mouse B cells. Nik-binding protein (NIBP), Ras-GAP SH3 binding protein 2 (G3BP1) and Trk-fused gene (Tfg) were identified by peptide mass fingerprinting in immunopurified CARMA1 complexes. The interaction of Tfg and CARMA1 was confirmed by co-immunoprecipitation and Blue native polyacrylamide gel electrophoresis using the anti CARMA1 and newly generated anti Tfg antibodies. This analysis revealed that CARMA1 formed complexes of 600-1000 kDa. Additionally, Tfg was found in complexes of 500-600 kDa which increased in size to ∼740 kDa upon overexpresssion.

## Introduction

The B cell receptor (BCR) is a membrane bound immunoglobulin (Ig) molecule consisting of two heavy chains (HC) of the μ type (μHC) and two light chains (LC) with variable and constant regions (Reth, 1991). The signaling unit Igα/β connects the BCR to Src, Syk and Bruton’s tyrosine kinases that initiate Ca^2+^ flux via SLP-65/BLNK (Wollscheid et al., 1999). Proximal signals recruit protein kinase Cβ (PKCβ) to the membrane, resulting in activation of downstream transcription factors, such as nuclear factor κB (NF-κB) (Hombach, 1990; Reth, 1991; DeFranco, 1992; Niiro, 2003; Thome, 2010). Classical NF-κB activation further downstream is achieved by proteasomal degradation of the Inhibitor of NF-κB (IκB) after its phosphorylation and ubiquitinylation, allowing translocation of p65/RelA-p50 or c-Rel/p50 heterodimers into the nucleus (Ghosh and Baltimore, 1990; May and Ghosh, 1997). This phosphorylation is a result of IκB-kinase (IKK) α and β activation via IKKγ/NEMO. An essential adaptor protein linking proximal BCR and T cell receptor (TCR) signaling to activation of the IKK complex, Jun N-terminal kinase (JNK), mTORC1 (mammamlian target of Rapamycin complex 1) and the sodium/glutamine antiporter ASCT2 (encoded by the *slc1a15* gene) is CARMA1/CARD11 (Egawa et al., 2003; Hara et al., 2003; Jun et al., 2003). CARMA1 consists of a CARD-domain, followed by a coiled-coil domain, a linker region, a PDZ and a SH3 domain and a guanylate kinase (GUK)-domain. CARMA1 forms a complex with the CARD containing B cell lymphoma (Bcl) 10 and Mucosa associated lymphoid tissue (MALT) 1, the CBM complex. Hence, null mutations of Bcl10 and MALT1 abrogate antigen receptor induced activation of NF-κB, JNK and mTORC1 (reviewed in Lu et al., 2018). CBM activation in B cells is achieved by phosphorylation of CARMA1 in its linker region by PKCβ, leading to opening of CARMA1. CARMA1 and Bcl10 interact through their CARD domains. The paracaspase domain of MALT1 interacts with the coiled-coil domain of CARMA1 and the IgG domains of MALT1 bind the Serine/Threonine rich region of Bcl10 (reviewed in Lu et al., 2018; Rawlings et al., 2006). Antigenic stimulation elicits higher order structures of the CBM complex by virtue of filaments formed by Bcl10. This creates an amplification loop, initiating MALT1 protease activaty as well as canonical NF-κB and JNK signaling. Moreover, MALT1 feeds back on the CBM complex through its protease activity. Further phosphorylation and ubiquitination of BcL10 induces more feedback and feed-forward events within the CBM complex. Thus, BCL10 is a positive and negative integrator of processes that tune the function and destruction of CBM complexes (Gehring et al., 2018).

While the CBM complex is required for humoral immunity it is also involved in diffuse large B cell lymphomagenesis through somatic gain of function mutations in CARMA1 (Lenz et al., 2008). In contrast, mutations in the genes encoding the CBM complex cause a variety of inborn immunodeficiencies, the “CBM-opathies” (Lu et al., 2018). The CBM mediated signaling pathways towards NF-κB and JNK are complex but quite well described. How CARMA1 is linked to mTORC1 controlled protein synthesis is not clear but Bcl10 is not required for this signaling branch (Hamilton et al., 2014; Lu et al., 2018). We hypothesized that previously not identified proteins interact with CARMA1 in B cells. Here we describe the enrichment of endogenous CARMA1 complexes through antibody affinity chromatography and the identification of three proteins co-purified with CARMA1, namely NIBP1, G3BP1 and Tfg. The putative interaction of CARMA1 and Tfg was confirmed by co-immunoprecipitation and Blue Native PAGE.

## Results and Discussion

The aim of this project was to identify new components of the lymphocyte specific CARMA1 signalling complex and therefore we chose to purify the endogenous CARMA1 complex through immuno-affinity purification from mouse B lymphoma cells. We generated two rabbit anti-CARMA1 peptide antibodies (584 and 947), directed against peptides outside the known CARMA1 domains. The affinity-purified antibodies detect murine full-length CARMA1 ectopically expressed in HeLa cells (Fig. 1a) as well as degradation product of CARMA1 likely resulting from overexpression (Fig. 1a). Specificity and suitability of the antibodies was confirmed by immunoprecipitation from 2 B cell lines, CH27 and WEHI231 (Fig. 1b), followed by excision of the putative CARMA1 band and mass spectrometric confirmation of its identity (Table 1).

**Table 1:**
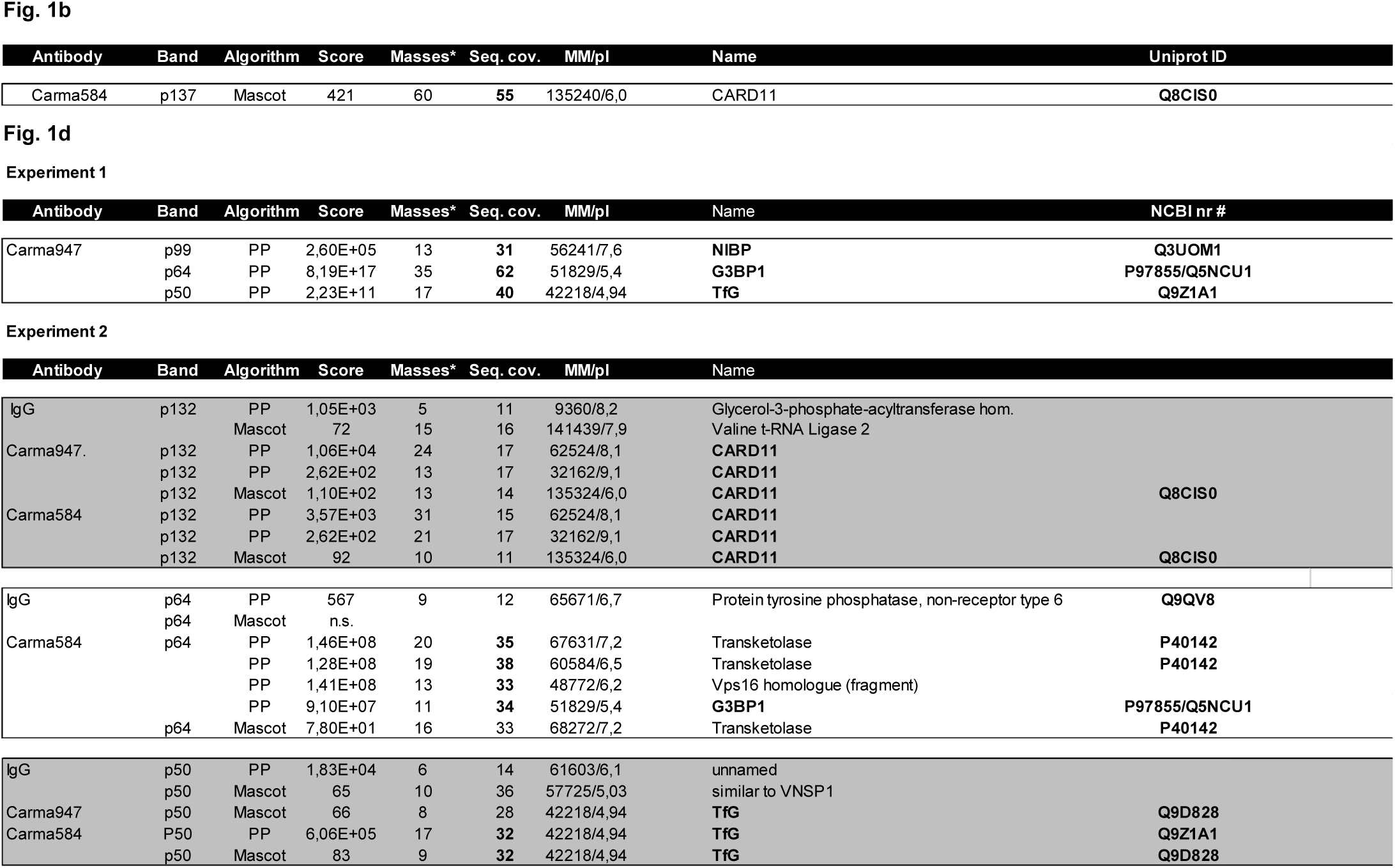
Summary of protein identification data. Specific bands were cut out, digested with Trypsin and the resulting peptides were eluted from the gel matrix. Peptides were then subjected to MALDI-TOF and masses were searched against current versions of NCBI nr database with two algorithms: MSFit (http://prospector.ucsf.edu/cgi-bin/msform.cgi?form=msfitstandard) and MASCOT (http://www.matrixscience.com/cgi/search_form.pl?FORMVER=2&SEARCH=PMF). Scores were considered to be significant in the case of MASCOT when indicated in the output window and for MSFit when a sequence coverage of > = 30% was achieved (with the exception of the data for Experiment 2, p132, since significant results were obtained in parallel with MASCOT). Masses refer to matched masses*. Seq. cov.: sequence coverage, MM/pI: molecular mass/isoelectric point. Uniprot ID: protein identification number (http://www.uniprot.org/). Proteins repeatedly identified are in bold.

**Figure 1.**
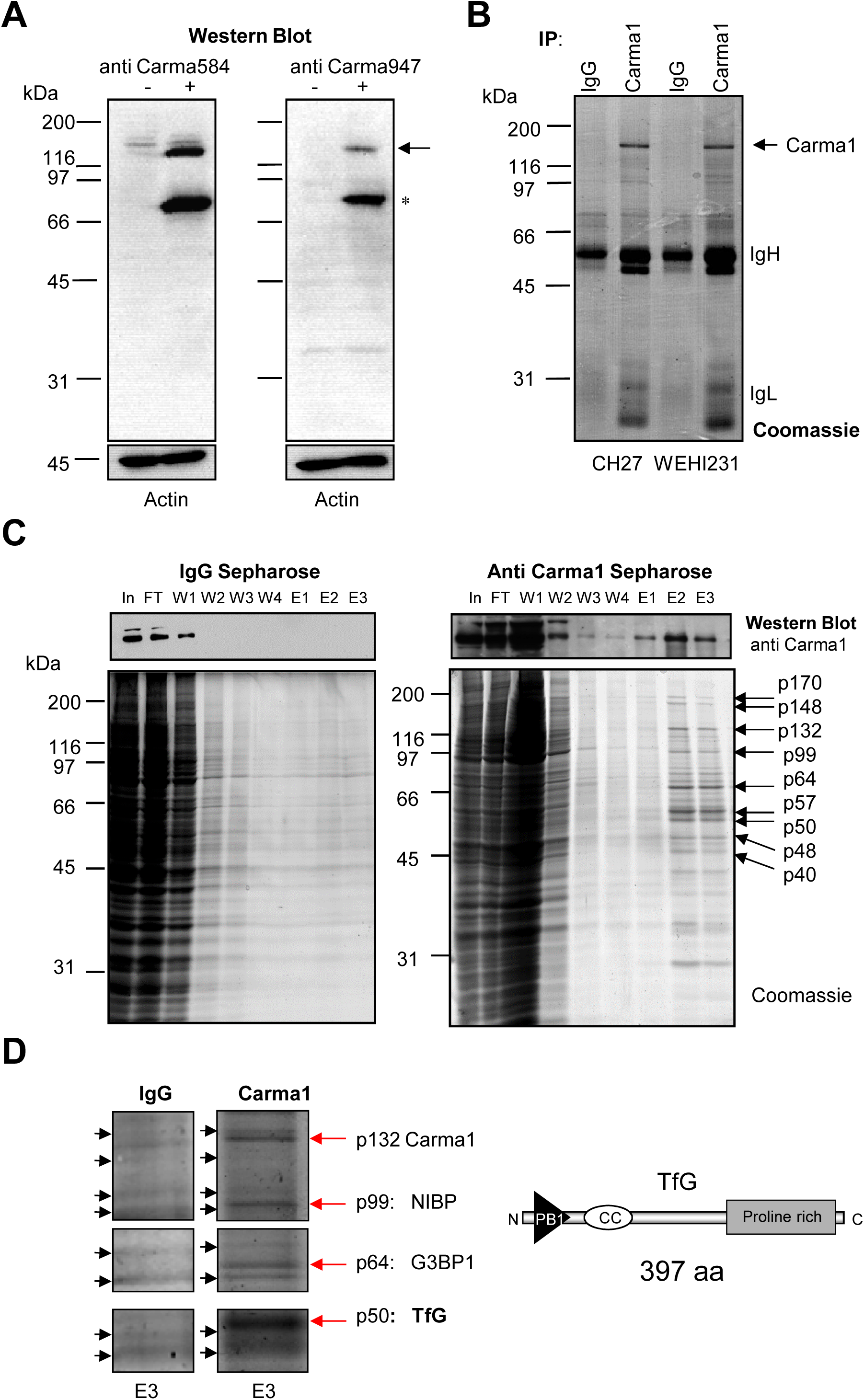
Identification of CARMA1 signalosome components. A) HeLa cells were left untransfected or transfected with a CARMA1 expression plasmid, lysed and analyzed by Western Blot with anti-CARMA1 antibodies 584 and 947, and with anti-actin antibodies. The arrow marks full-length CARMA1, * points to a CARMA1 degradation product in HeLa cells. Molecular mass standards (kDa) are indicated on the left. B) WEHI231 and CH27.LX B cells were lysed and lysates were subjected to immunoprecipitation with Protein A-purified anti-CARMA1 pre-immune serum (Pre) or with anti-CARMA1 antibody 584. Immunoprecipitates were subjected to 10 % SDS-PAGE, followed by colloidal Coomassie staining, and the band corresponding to CARMA1 (arrow) was cut out. IgH points to the heavy and light chains of the precipitating antibodies. Molecular mass standards (kDa) are indicated on the left. C) CH27.LX cells were lysed and lysates were subjected to immunoaffinity-purification with Protein A-purified anti-CARMA1 pre-immune serum (IgG), or anti-CARMA1 antibody 947 coupled covalently to Sepharose CL4B. Aliquots of input, flowthrough (FT), wash fractions (W) and eluate fractions (E) were analyzed by Western-Blotting with anti-CARMA1 antibodies (upper panel) or colloidal coomassie staining. D) Eluate fraction E3 of the IgG control Sepharose or the anti-CARMA1 Sepharose was separated by 10% SDS-PAGE and stained with colloidal Coomassie. Bands that appeared specifically in the eluate fractions of anti-CARMA1 Sepharose are marked with an arrow and their molecular masses are indicated (kDa). E) Drawn-to-scale schematic showing the predicted secondary structure of Tfg. N, N-terminus, C, C-terminus, PB1: Phox and Bem homology domain 1, CC: coiled-coiled domain.

Affinity-purified antibodies 584 and 947 as well as Protein A-purified CARMA1-947-preimmune serum were used to immunopurify CARMA1 from lysates of non-activated CH27.LX B cells via antibody affinity chromatography. Western Blot analysis of aliquots of input (In), flowthrough (FT), wash (W) and eluate (E) fractions (5% of each fraction demonstrated that CARMA1 eluted predominantly in fractions E2 and E3 (representative experiment shown in Fig. 1c, upper panel). Aliquots of input and flowthrough (5% of each fraction) and wash fractions along with eluate fractions (90% of the fractions) were further separated by 10% SDS-PAGE and stained with colloidal coomassie blue (representative experiment shown in Fig. 1c, lower panel). This experiment was performed two times with anti CARMA1-947 Sepharose and once with anti CARMA1-584-Sepharose in parallel with IgG-Sepharose (Table 1). Specific bands compared to the control eluate were cut out and identified by mass spectrometry (Table 1), revealing CARMA1 itself (p132), the IKK and NIK-binding protein (NIBP; p99), RasGAP-SH3-binding protein 2 (G3BP1, p64) as well as the Trk-fused-gene (p50, Tfg) (specific representative bands are marked in Figure 1d with red arrows, black arrows mark unspecific bands). Only proteins identified independently at least twice with significant cores were considered to be relevant and are marked in bold. Repetition of this experiment with anti BCR (45 min) activated B cells resulted in too many unspecific bands (data not shown). Surprisingly, we did not identify Bcl10 or MALT1 by mass spectrometry in eluates of the anti CARMA1 Sepharose, likely because we focused on distinctive bands. Bcl10 and MALT1 may have been covered by unspecific bands. NIBP has been shown to support canonical NF-κB and JNK activity (Hu et al., 2005; Qin et al., 2017), G3BP-1 is a Ras-GTPase-activating protein (SH3 domain)-binding protein (also known as Rasputin) belonging to a family of RNA binding proteins that regulate gene expression in response to environmental stresses by controlling mRNA stability and translation (Alam and Kennedy, 2019). Tfg consists of 397 amino acids structured into an N-terminal PB1 protein-protein-interaction domain (Sumimoto et al., 2007), a coiled-coil domain and a C-terminal Proline rich domain (Figure 1d). Tfg has been shown to interact with IKKγ (Miranda et al., 2006), promote cell size and oppose apoptosis antagonistic to ced-4/APAF-1 (Chen et al., 2008), be important for organization of the endoplasmic reticulum (ER; Johnson et al., 2015) and control organization of the ER-Golgi-intermediate compartment (ERGIC; Hanna et al., 2017). Moreover, Tfg is also involved in the ubiquitin-proteasome system (UPS; Yagi et al., 2014) and fusions of the *tfg* gene with other genes can lead to oncogenic fusion proteins that drive lymphomas and leukemias (Chase et al., 2010; Chong et al., 2018; Hernandez et al., 2002; Hernandez et al., 1999; Kim et al., 2016; Witte et al., 2011). The intrinsic oligomerization activity of Tfg (Johnson et al., 2015), its interaction with the IKK complex, its positive effect on cell growth and its structural function in the ER/ERGIC suggested that it could be a functional constituent of the CBM complex, connecting it to IKK or mTORC1 enhanced mRNA translation. Thus, we focused on Tfg.

To test further for an interaction between CARMA1 and Tfg we used CH12 B cells (Bishop and Haughton, 1986) to overexpress either N-terminally (HA-Tfg) or C-terminally (Tfg-HA) HA-tagged Tfg. Conversely, we silenced Tfg expression in CH12 B cells by stable shRNA-interference and we generated specific, affinity-purified anti-Tfg peptide antibodies (Figure 2a). Using these antibodies we confirmed the presence of Tfg in eluates of the anti CARMA1 584 column by Western Blotting (Figure 2b). To corroborate the CARMA1-Tfg interaction we immunoprecipitated Tfg from lysates of Tfg-silenced, control or Tfg-overexpressing CH12 cells using our own anti Tfg antiserum (Figure 2c). While being present at comparable amounts in the total cell lysate, CARMA1 co-precipitated dose-dependently with Tfg regardless of short term BCR or CD40 stimulation (Figure 2c). Long term stimulation may alter the stoichiometry of the Tfg-CARMA1 complex, a possibility that we have not tested here. We noted that still tiny amounts of Tfg as well as CARMA1 were precipitated from CH12 cells with silenced Tfg expression, suggesting that silencing was not 100%. We failed to co-immunoprecipitate CARMA1 with HA-tagged Tfg using anti-HA antibodies in several attempts (data not shown). This may be due to sterical hindrance. Moreover, in our column-based immuno-affinity purification approach the protein concentration in the cell lysate and the avidity of the anti CARMA1 antibody avidity on the affinity matrix were higher than under IP conditions, likely contributing to stabilization of CARMA1 complexes. Lastly, protease activity towards CARMA1 may be associated with Tfg under these conditions, which could be addressed by specific inhibition.

**Figure 2.**
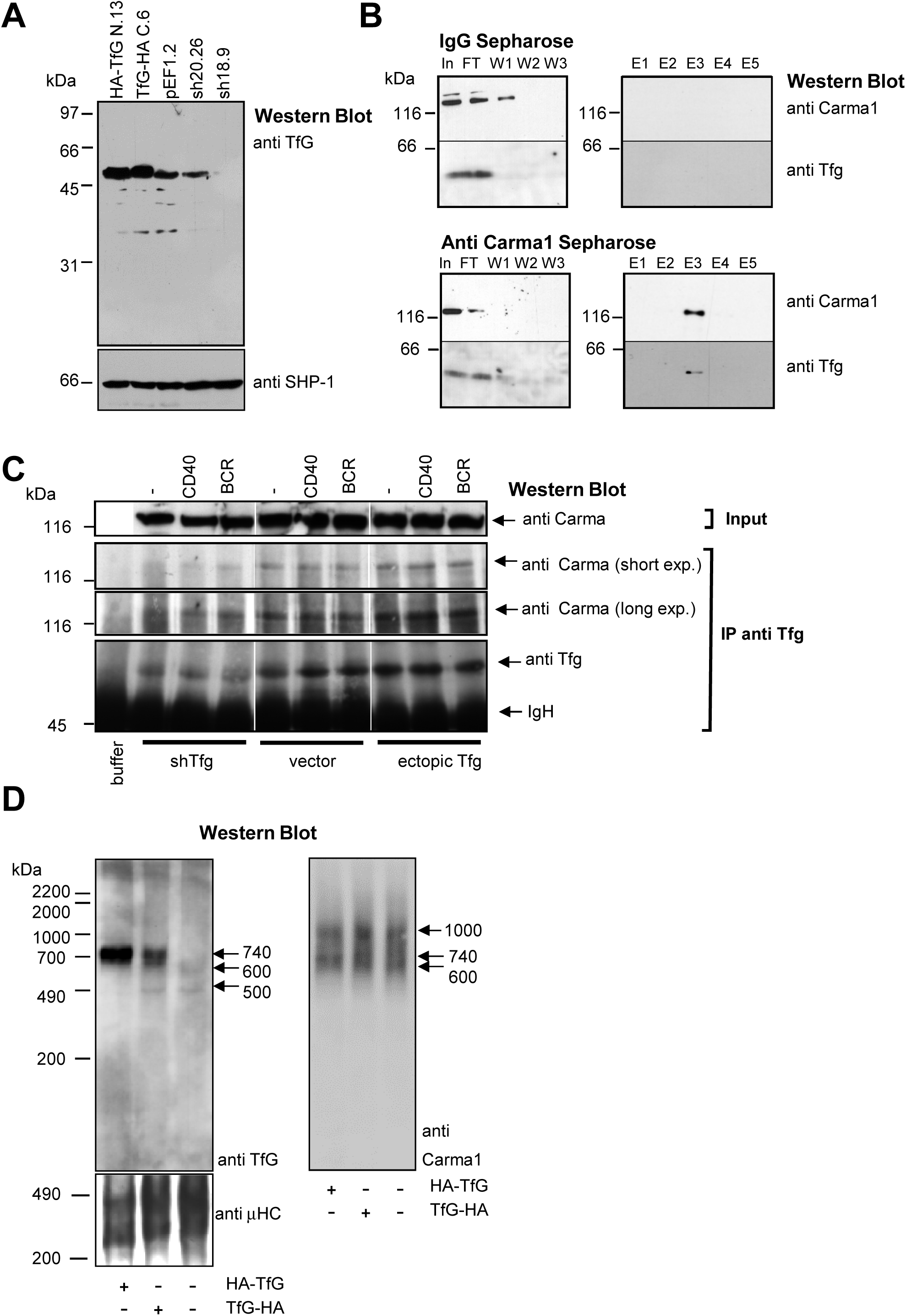
Analysis of CARMA1 and Tfg complexes by Co-immunoprecipitation and Blue Native PAGE. A) CH12 cells expressing a Tfg-specific shRNA (clone 18.9), a control shRNA (clone 20.26), an empty vector (pEF1.3) or ectopic TfG with a C- or N-terminal HA-Tag (Tfg-HA, clone C.6, HA-Tfg, N.13) were generated by transfection and single cell cloning. Tfg expression was assessed by Western Blotting of 10% SDS-PAGE gels as indicated. Molecular mass standards (kDa) are shown on the left. B) CH12 cells were lysed and lysates were subjected to immunoaffinity-purification with Protein A-purified anti-CARMA1 pre-immune serum (IgG), or anti-CARMA1 antibody 584 coupled covalently to Sepharose CL4B. Aliquots of input, flowthrough (FT), wash fractions (W) and eluate fractions (E) were analyzed by Western-Blotting with anti-CARMA1 584 and anti-Tfg antibodies. Molecular mass standards (kDa) are shown on the left. C) CH12 cells expressing a Tfg-specific shRNA (clone 18.9), an empty vector (pEF1.3) or ectopic TfG with a C-terminal HA-Tag (Tfg-HA, clone C.6) were left unstimulated or stimulated with agonistic anti-BCR and anti-CD40 mAbs for 3 min. Cells were lysed, an aliquot was subjected to 6% SDS-PAGE and analyzed directly by Western-Blotting with anti-CARMA1 584 (input, top panel). The remaining lysate was subjected to immunoprecipitation with rabbit anti-Tfg antibody and a control with buffer only was included. IPs were then analyzed by Western-Blotting with anti-CARMA1 584 and anti-Tfg antibodies. For CARMA1, two exposure times (short exp. and long exp.) are shown. Molecular mass standards are indicated on the left (kDa). Within one panel, all cutouts are from the same film and have the same exposure time. D) Lysates of CH12 cells expressing an empty vector (clone pEF1.3) or ectopic TfG with an C- or N-terminal (HA-Tfg, clone N.13, Tfg-HA, clone C.6) were subjected to BN-PAGE, transferred onto a PVDF membrane and the membrane was cut into four stripes that were incubated with anti-Tfg antibody, anti-CARMA1 584 antibody or anti-BCR μ heavy chain (μHC) antibody. Molecular mass standards (bovine heart mitochondria) are indicated on the left (kDa).

To further confirm the existence of a CARMA1-Tfg complex, avoid possible sterical hindrance and determine the size of the CARMA1-Tfg complex, we separated lysates of CH12 cells with endogenous Tfg expression (empty vector) and CH12 clones expressing N-or C-terminally HA-tagged Tfg by Blue native polyacrylamide gel electrophoresis (BN-PAGE; Schägger and von Jagow, 1991) (Fig. 2d). To avoid dialysis of the whole cell lysates (Camacho-Carvajal et al., 2004) we developed a simple alternative technique to prepare cell lysates for BN-PAGE (see Materials & Methods). To detect Tfg and CARMA1 by BN-PAGE, whole cell lysates were separated twice in parallel and blotted. The resulting Blot was cut into stripes and probed separately with antibodies against Tfg and CARMA1 (Figure 2c). To demonstrate equal loading, the blot probed with anti Tfg antibody was afterwards probed with a directly horse radish peroxidase conjugated anti BCR μ heavy chain antibody (Figure 2c). This experiment revealed that the Tfg complex has a molecular mass of 500-600 kDa in B cells, which increases up to 740 kDa upon Tfg overexpression. These data are in accord with previous reports showing octameric oligomers of Tfg (Johnson et al., 2015). We suggest that in B cells, Tfg is part of a CARMA1 subcomplex of about 600 kDa and that Tfg overexpression induces a 740 kDa complex containing at least CARMA1 and Tfg. In summary, we provide three lines of evidence for an interaction of CARMA1 with Tfg in murine B lymphoma cells: First, identification of Tfg by mass spectrometry and Western Blotting in eluates of anti-CARMA1 Sepharose, second, co-purification of CARMA1 with Tfg in anti-Tfg immunoprecipitates and third, a concomitant shift towards higher molecular mass of Tfg and CARMA1 complexes upon Tfg overexpression. It remains to be determined whether there is a direct and functional interaction between Tfg and CARMA1. Due to linkage of the CBM complex to the mTORC1 pathway and the supportive role of Tfg in regulation of cell size and ER function it is tempting to speculate that Tfg is involved in these processes downstream of the BCR via the CBM complex.

## Abbreviations

Ag: antigen
Bcl10: B cell lymphoma 10
BCR: B cell receptor
Btk: Bruton’s tyrosine kinase
ER: endoplasmic reticulum
ERGIC: ER Golgi intermediate compartment
Ig: immunoglobulin
HC: heavy chain
LC: light chain
IκB: Inhibitor of NF-κB
IKK: IκB-kinase
LC: light chain
JNK: Jun N-terminal kinase
MALT1: Mucosa-associated lymphoid tissue lymphoma translocation protein 1
MALDI-TOF: matrix assisted laser desorption ionization time-of-flight
mTORC1: mammalian target of Rapamycin complex 1
PKCβ: protein kinase Cβ
NF-κB: nuclear factor κB
TCR: T cell receptor
UPS: ubiquitin-proteasome system.

## Acknowledgements

The current work was supported by BMBF/IZKF grant A7 (Universitätsklinikum Erlangen, to D.M. and H.M.J), and the Deutsche Forschungsgemeinschaft (DFG; GK1660 and TRR130, to D.M. and H.M.J.). We thankfully acknowledge the group of Prof. Dr. Michael Karas, Instrumental Analytical Chemistry, Institute of Pharmaceutical Chemistry, University of Frankfurt, Germany, for performing mass spectrometric analyses.

## Materials and Methods

### Transfection

HeLa cells were transfected with full length murine CARMA1 cDNA cloned into pCMVSport6 (obtained from RZPD, Berlin, Germany, clone ID IRAK0961H0B208Q2) with Lipofectamine (Thermo Fisher) according to standard procedures. CH12 cells (5 × 10^6^) were electroporated with 10μg of cDNA by nucleofection (Solution “R”, program “K-005”, Amaxa). Single clones were obtained by limiting dilution of cell kept in selection medium (G418, 400 μg/ml). N- or C-terminally HA-tagged Tfg cDNAs were generated by PCR using a full length cDNA clone (obtained by RZPD, Berlin, Germany, clone ID IRAKp961H03208Q2) as template with the following primers containing BamHI or NotI restriction sites: Tfg_HA_BamHI_for, 5’GATCGGATCCGCCACCATGTACCCATACGATGTTCCAGATTACGCTAA, Tfg_NotI_back, 5’CTAGGCGGCCGCTTATCGATAACCAGGTCCAGGT, Tfg_BamHI_for, 5’ GATCGGATCCGCCACCATGAACGGACAGTTGGACCTAAG, Tfg_HA_NotI_back, 5’, CTAGGCGGCCGCTTAAGCGTAATCTGGAACATCGTATGGGTATCGATAACCAGGTCCAGGT. PCR products were cloned into pEF1 (Invitrogen) and identity was confirmed by sequencing. Vector driven anti-Tfg shRNA expression was achieved using transfection of commercially available vectors based on the lentiviral pLKO backbone (Openbiosystems, Huntsville, AL), followed by single cell cloning and Puromycin selection (0.4μg/ml). Of five constructs obtained, one silenced Tfg efficiently (#N00000079118, Clone ID NM_019678.1-1680s1c1, targeting the 3’ UTR). Clones derived from this construct were named 18.xx. Another construct (#N0000079120, Clone ID NM_019678.1-551s1c1, targeting the coding sequence) was therefore used as control. Clones derived from this construct were named 20.xx.

### Immunoprecipitation

20 × 10^6^ CH27 (Bishop and Haughton, 1986) or WEHI231 (Boyd et al., 1981) cells were lysed in 1 ml of IP lysis buffer (1% NP40, 25 mM HEPES/KOH pH 7.5, 120 mM NaCl, 30 mM NaF, 5 mM ethylene-diamine-tatra-acetate (EDTA), 5 mM 6-aminocaproic acid, 1 mM phenyl-methyl-sulfonyl-fluoride (PMSF), 1 mM N-ethyl-maleimide (NEM), 1 mM Sodium Vanadate) for 10 min on ice and centrifuged for 5 min at 5000 g at 4°C After determination of the protein concentration by the BCA test (Pierce, Rockford, IL), 2-5 μg of antibody and 30 μl of equilibrated Protein A- or Protein-G-Sepharose (Pierce, Rockford, IL) were added and the mixture was rocked head over head for 1.5 h at 4°C Beads were washed 2 times by repeated centrifugation (1000g, 1min), resuspension in ice-cold lysis buffer and aspiration of the supernatants, boiled for 5 min in SDS-sample loading buffer and applied to SDS-PAGE.

### Blue Native (BN) polyacrylamide gel electrophoresis (PAGE)

BN-PAGE was essentially performed according to previously published procedures (Schägger and von Jagow, 1991) but cell lysates were prepared as follows to avoid dialysis: 40 × 10^6^ cells were lysed in 1 ml of buffer containing 1% NP40, 120 mM NaCl, 30 mM NaF, 25 mM Imidazole pH 7.6, 5 mM EDTA, 5 mM 6-aminocaproic acid, 1 mM PMSF, 1 mM NEM, 1 mM Vanadate) for 10 min on ice and centrifuged for 5 min at 5000g. Supernatants were immediately mixed with an equal volume of ice-cold 85% Sucrose in 120 mM NaCl, 30 mM NaF, 25 mM Imidazole pH 7.6. Then, samples were diluted 1:2 with ice-cold 1M 6-aminocaproic acid to achieve final concentrations of 2.25 % sucrose and 500 mM 6-aminocaproic acid. Samples were kept on ice and shortly before electrophoresis, 2.5 % of a 5% Coomassie G-250 stock solution in 500 mM 6-aminocaproic acid were added. Electrophoresis was performed with non-linear 3-13 % acrylamide gels containing 25 mM Imidazole, pH 7.5 and 500 mM 6-aminocaproic acid. Cathode buffer was 50 mM Tricine, 7.5 mM Imidazole, 0.02 % Coomassie G250 for the first third of the gel and then changed to 50 mM Tricine, 7.5 mM Imidazole, 0.002 % Coomassie G250. Anode buffer was 25 mM imidazole/HCl pH 7.0 and gels were run at 4°C with 200 μg of bovine heart mitochondria as molecular mass standards (kind gift of Dr. Herrmann Schägger, University of Frankfurt, Germany). Proteins were transferred to PVDF membranes for 45 min at 400 mA, membranes were destained with MeOH in water and blocked in 5 % skim milk powder in 25 mM Tris/HCl pH7.5, 150 mM NaCl, 0.1% Tween-20 (TBST).

### Immunoaffinity purification

1-3 × 10^8^ CH12 or CH27.LX cells (Bishop and Haughton, 1986) were lysed in 1 ml of 1% NP40, 120 mM NaCl, 30 mM NaF, 25 mM Tris/HCl pH 7.4, 5 mM EDTA, 5 mM 6-aminocaproic acid, 1 mM PMSF, 1 mM NEM, 1 mM Vanadate) for 10 min on ice and centrifuged for 5 min at 5000g. Supernatants were immediately loaded on anti-CARMA1 affinity columns or rabbit IgG affinity columns (Sepharose CL-4B, GE healthcare, ∼ 1 mg antibody/ml bedvolume) equivalent to 200 μg antibody and allowed to pass via gravity flow (∼ 5-10 min per ml) in the cold-room. Columns were washed with 3 × 1 ml of lysis buffer and eluted with 5 × 100 μl of 0.2 M glycine, 0.1 % SDS, Ph 2.5 in the cold-room. Aliquots were removed for Western Blotting and input, flow-through, wash and elution fractions were analyzed by 10 % SDS-PAGE and colloidal Coomassie staining. Briefly, gels were fixed in 40% EtOH, 10% acetic acid for 60 min, washed with distilled water for 2 × 10 min and the gel was soaked in staining solution (20% MeOH, 80% Coomassie solution [0.1% Coomassie Brilliant Blue G-250, 2% ortho-phosphoric-acid, 10% ammonium sulfate]) overnight. The gels were then soaked several times in 1% acetic acid until background was clear. Bands of interest were cut out in parallel with bands of control gels. Bands were digested with trypsin, peptides were eluted and analyzed by matrix assisted laser desorption ionization time-of-flight (MALDI-TOF) mass spectrometry as described previously (Mielenz et al., 2005; Vettermann et al., 2011). Masses were searched against current versions of NCBI nr databases with MSFit (http://prospector.ucsf.edu/cgi-bin/msform.cgi?form=msfitstandard) and MASCOT (http://www.matrixscience.com/cgi/search_form.pl?FORMVER=2&SEARCH=PMF).

### Antibodies

Anti-CARMA1 antibodies 584 and 947 and anti-Tfg antibody were generated in rabbits against synthetic peptides (CARMA1 peptide 584-979, EDAPHRSTVEEDND, CARMA1 peptide 947-963, TEKHEELDPENELSRN, Tfg peptide 132-145 STSIPENDTVDGRE), whereby numbers indicate the first amino acid in the peptide according to the numbering Uniprot ID Q8CISO (CARMA1) or Q9Z1A1 (Tfg). Peptides contained an N-terminal cysteine residue and were coupled to keyhole-limpet-hemocyanin (KLH) using a sulfhydryl-directed bifunctional cross-linker. New Zealand white rabbits were immunized with KLH-coupled peptides according to standard procedures (Harlow, 1988) (http://www.pineda-abservice.de/). For antibody affinity purification, peptides were coupled via the cysteines to Sulfo-link resine (Pierce, Rockford, IL) according to the manufacturer’s instructions. Antibody purification on peptide columns was performed according to standard procedures (Harlow, 1988). Affinity-purified antibodies were used at 0.5-1 μg/ml for Western-Blotting. Rat anti-CD40 and anti-B cell receptor antibodies were purified from the IC10 and b.7.6 hybridomas (Julius et al., 1984; Santos-Argumedo et al., 1994).

### SDS-PAGE and Western Blotting

Cells were washed in PBS and lysed in IP lysis buffer (see above). Western Blotting of 12×15cm^2^ SDS-PAGE gels was performed according to standard procedures. Membranes were blocked in 5% skim milk powder in TBST and washed in TBST. Primary antibodies were diluted in 3% BSA in TBST containing 0.02%NaN3, secondary antibodies were diluted in 5% skim milk powder in TBST. Blots were developed by enhanced chemiluminescence. Antibodies were rabbit Ig anti-CARMA1 and anti-Tfg (described above), rabbit Ig anti-mouse Actin (Sigma-Aldrich), horse radish peroxidase conjugated goat Ig anti-mouse IgG (H+L; Biorad), horse radish peroxidase conjugated goat Ig anti-mouse Igμ (Southern Biotech).

## References

Alam, U., and Kennedy, D. (2019). Rasputin a decade on and more promiscuous than ever? A review of G3BPs. Biochim Biophys Acta Mol Cell Res 1866, 360–370.

Bishop, G.A., and Haughton, G. (1986). Induced differentiation of a transformed clone of Ly-1+ B cells by clonal T cells and antigen. Proc Natl Acad Sci U S A 83, 7410–7414.

Boyd, A. W., J. W. Goding, and J. W. Schrader. (1981). The regulation of growthand differentiation of a murine B cell lymphoma. I. Lipopolysaccharide-induceddifferentiation. J. Immunol. 126, 2461

Camacho-Carvajal, M.M., Wollscheid, B., Aebersold, R., Steimle, V., and Schamel, W.W. (2004). Two-dimensional Blue native/SDS gel electrophoresis of multi-protein complexes from whole cellular lysates: a proteomics approach. Mol Cell Proteomics 3, 176–182.

Chase, A., Ernst, T., Fiebig, A., Collins, A., Grand, F., Erben, P., Reiter, A., Schreiber, S., and Cross, N.C. (2010). TFG, a target of chromosome translocations in lymphoma and soft tissue tumors, fuses to GPR128 in healthy individuals. Haematologica 95, 20–26.

Chen, L., McCloskey, T., Joshi, P.M., and Rothman, J.H. (2008). ced-4 and proto-oncogene tfg-1 antagonistically regulate cell size and apoptosis in C. elegans. Curr Biol 18, 1025–1033.

Chong, M.L., Cheng, H., Xu, P., You, H., Wang, M., Wang, L., and Ho, H.H. (2018). TFG-RARA: A novel fusion gene in acute promyelocytic leukemia that is responsive to all-trans retinoic acid. Leuk Res 74, 51–54.

Egawa, T., Albrecht, B., Favier, B., Sunshine, M.J., Mirchandani, K., O’Brien, W., Thome, M., and Littman, D.R. (2003). Requirement for CARMA1 in antigen receptor-induced NF-kappa B activation and lymphocyte proliferation. Curr Biol 13, 1252–1258.

Gehring, T., Seeholzer, T., and Krappmann, D. (2018). BCL10 - Bridging CARDs to Immune Activation. Front Immunol 9, 1539.

Ghosh, S., and Baltimore, D. (1990). Activation in vitro of NF-kappa B by phosphorylation of its inhibitor I kappa B. Nature 344, 678–682.

Hamilton, K.S., Phong, B., Corey, C., Cheng, J., Gorentla, B., Zhong, X., Shiva, S. Kane, L.P. (2014). T cell receptor-dependent activation of mTOR signaling in T cells is mediated by CARMA1 and MALT1, but not Bcl10. Sci Signal. 2014 Jun 10;7(329):ra55. doi: 10.1126/scisignal.2005169.

Hanna, M.G.t., Block, S., Frankel, E.B., Hou, F., Johnson, A., Yuan, L., Knight, G., Moresco, J.J., Yates, J.R., 3rd, Ashton, R., et al. (2017). TFG facilitates outer coat disassembly on COPII transport carriers to promote tethering and fusion with ER-Golgi intermediate compartments. Proc Natl Acad Sci U S A 114, E7707–E7716.

Hara, H., Wada, T., Bakal, C., Kozieradzki, I., Suzuki, S., Suzuki, N., Nghiem, M., Griffiths, E.K., Krawczyk, C., Bauer, B., et al. (2003). The MAGUK family protein CARD11 is essential for lymphocyte activation. Immunity 18, 763–775.

Harlow, E.L. David (1988). Antibodies: A Laboratory Manual (Cold Spring Harbour: Cold Spring Harbour Press).

Hernandez, L., Bea, S., Bellosillo, B., Pinyol, M., Falini, B., Carbone, A., Ott, G., Rosenwald, A., Fernandez, A., Pulford, K., et al. (2002). Diversity of genomic breakpoints in TFG-ALK translocations in anaplastic large cell lymphomas: identification of a new TFG-ALK(XL) chimeric gene with transforming activity. Am J Pathol 160, 1487–1494.

Hernandez, L., Pinyol, M., Hernandez, S., Bea, S., Pulford, K., Rosenwald, A., Lamant, L., Falini, B., Ott, G., Mason, D.Y., et al. (1999). TRK-fused gene (TFG) is a new partner of ALK in anaplastic large cell lymphoma producing two structurally different TFG-ALK translocations. Blood 94, 3265–3268.

Hombach, J., Tsubata, T., Leclercq, L., Stappert, H., and Reth, M. (1990). Molecular components of the B-cell antigen receptor complex of the IgM class. Nature 343, 760–762.

Hu, W.H., Pendergast, J.S., Mo, X.M., Brambilla, R., Bracchi-Ricard, V., Li, F., Walters, W.M., Blits, B., He, L., Schaal, S.M., et al. (2005). NIBP, a novel NIK and IKK(beta)-binding protein that enhances NF-(kappa)B activation. J Biol Chem 280, 29233–29241.

Johnson, A., Bhattacharya, N., Hanna, M., Pennington, J.G., Schuh, A.L., Wang, L., Otegui, M.S., Stagg, S.M., and Audhya, A. (2015). TFG clusters COPII-coated transport carriers and promotes early secretory pathway organization. EMBO J 34, 811–827.

Julius, M.H., Heusser, C.H., and Hartmann, K.U. (1984). Induction of resting B cells to DNA synthesis by soluble monoclonal anti-immunoglobulin. Eur J Immunol 14, 753–757.

Jun, J.E., Wilson, L.E., Vinuesa, C.G., Lesage, S., Blery, M., Miosge, L.A., Cook, M.C., Kucharska, E.M., Hara, H., Penninger, J.M., et al. (2003). Identifying the MAGUK protein Carma-1 as a central regulator of humoral immune responses and atopy by genome-wide mouse mutagenesis. Immunity 18, 751–762.

Kim, A.Y., Lim, B., Choi, J., and Kim, J. (2016). The TFG-TEC oncoprotein induces transcriptional activation of the human beta-enolase gene via chromatin modification of the promoter region. Mol Carcinog 55, 1411–1423.

Lenz, G., Davis, R.E., Ngo, V.N., Lam, L., George, T.C., Wright, G.W., Dave, S.S., Zhao, H., Xu, W., Rosenwald, A., et al. (2008). Oncogenic CARD11 mutations in human diffuse large B cell lymphoma. Science 319, 1676–1679.

Lu, H.Y., Bauman, B.M., Arjunaraja, S., Dorjbal, B., Milner, J.D., Snow, A.L., and Turvey, S.E. (2018). The CBM-opathies-A Rapidly Expanding Spectrum of Human Inborn Errors of Immunity Caused by Mutations in the CARD11-BCL10-MALT1 Complex. Front Immunol 9, 2078.

May, M.J., and Ghosh, S. (1997). Rel/NF-kappa B and I kappa B proteins: an overview. Semin Cancer Biol 8, 63–73.

Mielenz, D., Vettermann, C., Hampel, M., Lang, C., Avramidou, A., Karas, M., and Jäck, H.M. (2005). Lipid rafts associate with intracellular B cell receptors and exhibit a B cell stage-specific protein composition. J Immunol 174, 3508–3517.

Miranda, C., Roccato, E., Raho, G., Pagliardini, S., Pierotti, M.A., and Greco, A. (2006). The TFG protein, involved in oncogenic rearrangements, interacts with TANK and NEMO, two proteins involved in the NF-kappaB pathway. J Cell Physiol 208, 154–160.

Niiro, H., and Clark, E.A. (2003). Branches of the B cell antigen receptor pathway are directed by protein conduits Bam32 and CARMA1. Immunity 19, 637–640.

Qin, M., Zhang, J., Xu, C., Peng, P., Tan, L., Liu, S., and Huang, J. (2017). Knockdown of NIK and IKKbeta-Binding Protein (NIBP) Reduces Colorectal Cancer Metastasis through Down-Regulation of the Canonical NF-kappaBeta Signaling Pathway and Suppression of MAPK Signaling Mediated through ERK and JNK. PLoS One 12, e0170595.

Rawlings, D.J., Sommer, K., and Moreno-Garcia, M.E. (2006). The CARMA1 signalosome links the signalling machinery of adaptive and innate immunity in lymphocytes. Nat Rev Immunol 6, 799–812.

Reth, M. (1991). Signal transduction in B cells. Curr Opin Immunol 3, 340–344.

Santos-Argumedo, L., Gordon, J., Heath, A.W., and Howard, M. (1994). Antibodies to murine CD40 protect normal and malignant B cells from induced growth arrest. Cell Immunol 156, 272–285.

Schägger, H., and von Jagow, G. (1991). Blue native electrophoresis for isolation of membrane protein complexes in enzymatically active form. Anal Biochem 199, 223–231.

Sumimoto, H., Kamakura, S., and Ito, T. (2007). Structure and function of the PB1 domain, a protein interaction module conserved in animals, fungi, amoebas, and plants. Sci STKE 2007, re6

Thome M, Charton JE, Pelzer C, Hailfinger S. (2010). Antigen receptor signaling to NF-kappaB via CARMA1, BCL10, and MALT1. Cold Spring Harb Perspect Biol. 2010 Sep;2(9):a003004.

Vettermann, C., Castor, D., Mekker, A., Gerrits, B., Karas, M., Jäck, H.M. (2011). Proteome profiling suggests a pro-inflammatory role for plasma cells through release of high-mobility group box 1 protein. Proteomics. 11(7):1228–37

Witte, K., Schuh, A.L., Hegermann, J., Sarkeshik, A., Mayers, J.R., Schwarze, K., Yates, J.R., 3rd, Eimer, S., and Audhya, A. (2011). TFG-1 function in protein secretion and oncogenesis. Nat Cell Biol 13, 550–558.

Wollscheid, B., Wienands, J., and Reth, M. (1999). The adaptor protein SLP-65/BLNK controls the calcium response in activated B cells. Curr Top Microbiol Immunol 246, 283-288; discussion 288-289.

Yagi, T., Ito, D., and Suzuki, N. (2014). Evidence of TRK-Fused Gene (TFG1) function in the ubiquitin-proteasome system. Neurobiol Dis 66, 83–91.

